# Energetic costs of osmo- and ionoregulatory responses to desiccation and recovery in a model insect

**DOI:** 10.64898/2026.11.17.767398

**Authors:** Jacinta D. Kong, Susan M. Bertram, Heath A. MacMillan

## Abstract

Climate change is expected to increase the frequency, duration and intensity of weather events that cause desiccation stress to small-bodied terrestrial insects. Whether a desiccation event may cause mortality depends on the time-dependent osmo- and ionoregulation responses of insects to desiccation and their recovery. Yet these responses and their associated energetic costs are not always clear for a range of mesic species. Here, we characterised the ability of house crickets (*Acheta domesticus*) to physiologically resist and recover from desiccation. We measured water loss from different tissues, changes in haemolymph volume and in haemolymph ion concentration as well as stored macromolecules (lipids, carbohydrates and proteins) use over a time-course of desiccation (0, 3, 6 or 18 h) and recovery with access to water and with or without access to food (18 h desiccation and 6 h recovery). During 18 h of desiccation, crickets regulated haemolymph ion concentration, avoiding hyperkalaemia and hypernatraemia, despite losing up to 14.5 and 21.1% of their body water for female and males respectively, across all body compartments. Recovery represented a distinct physiological phase to acute stress with a restoration of ion concentrations and body water with significant energetic contributions from stored macromolecules. We conclude crickets employ a general two-phase physiological strategy to withstand and recover from a short-term acute desiccation stress, establishing a physiological baseline to evaluate potential desiccation stress, and providing key insight into the time-dependency of osmoregulation and ion regulation for insects encountering fluctuating desiccation stress under climate change.

## Introduction

Water is a major component of the abiotic environment that is essential to support terrestrial life (Hadley, 1994). Desiccation is a key abiotic stressor for small arthropods on land because their small body mass and high surface area to volume ratio can facilitate water loss to the environment when liquid water is scarce or when in arid habitats (O’Donnell, 2022). Terrestrial arthropods can resist or tolerate an acute desiccation stress using a variety of morphological, behavioural and physiological adaptations (Addo-Bediako et al., 2001; Hadley, 1994). These adaptations have allowed arthropods to successfully colonise dry habitats and overcome the biophysical limitations of their small size (Chown et al., 2011; Hadley, 1994; Halberg and Denholm, 2024; O’Donnell, 2022; Sinclair et al., 2024). However, climate change is expected to change the frequency, duration and intensity of extreme weather events, such as heat waves, or drought, increasing the duration of acute desiccation stress which, in turn, puts pressure on the ability of arthropods to maintain their water balance (Harvey et al., 2023; Johnson et al., 2026; Le Bivic et al., 2026). Potential climate change induced desiccation stress has been overlooked compared to thermal stress in the literature, yet desiccation affects insect activity, reproduction and survival, and intersects with thermoregulation and thermal tolerance through evaporative water loss (Botsch et al., 2024; Brown et al., 2023; Dahake et al., 2024; Simmons et al., 2023). How desiccation stress affects species under climate change then becomes a problem to understand the time-dependent nature of an individual’s response and recovery to acute desiccation stress as these responses over time will determine the effective severity of a desiccation event unfolding over increasing timescales for an individual.

An insect’s capacity to resist or tolerate bouts of desiccation depends on their response throughout the stress event and their recovery in benign environments. An insect progressively loses body water during acute desiccation stress and most species resist or tolerate these water losses until death (Folk and Bradley, 2003; Naidu, 2001a). Most insects show strong capacity for recovery from desiccation if provided fluids and/or food, including restoration of mass, haemolymph volume or ion concentration to baseline values (Albers and Bradley, 2004; Folk and Bradley, 2003; Hyatt and Marshall, 1977; Hyatt and Marshall, 1985; Naidu, 2001b; Spring et al., 1986). If a desiccation event is too severe, then an individual may not be able to fully recover and is potentially more vulnerable to the next desiccation event (Harvey et al., 2023). Thus, whether an individual can maintain homeostasis during desiccation stress and recover it afterward are both important for interpreting responses to desiccation on natural time scales.

Ionoregulation and osmoregulation are key physiological mediators of the desiccation response that allow individuals to tolerate water loss. These processes involve the renal system, cellular membranes compartmentalising various body components, the respiratory system, osmolytes and free cations, as well as stored macromolecules (Bosua et al., 2026; Gibbs et al., 2003; Halberg and Denholm, 2024; Machin, 1981). A reduction in haemolymph volume during desiccation, measured gravimetrically, indicates water is lost from the haemocoel in insects, but this method does not quantify water loss in other tissues (e.g. gut, cuticle, muscles etc.) that comprise total water loss (Folk and Bradley, 2003; Gefen and Ar, 2005; Naidu, 2001a). Haemolymph ion concentration and haemolymph volume are intrinsically linked, whereby ion concentrations (e.g. Na^+^ or K^+^ as primary cations) or total osmolality are expected to increase with a decrease in haemolymph volume (Albers and Bradley, 2004; Archer et al., 2007). If increases in ion concentrations continues during a desiccation stress as haemolymph volume declines, hyperkalaemia and/or hypernatraemia may occur, which disrupt cellular function and can lead to cellular injury, as described for other acute abiotic stressors (e.g. acute cold stress can co-occur with desiccation and also involves ionoregulatory collapse (MacMillan and Sinclair, 2011; Williams and Lee, 2011)). Yet, many studies indicate insects are actively ionoregulating during desiccation in ways that enhance their tolerance of water loss, resulting in lower-than-expected increases in haemolymph ion concentrations observed as haemolymph volume decreases, and demonstrating the importance of osmo- and ionoregulation in mediating the desiccation response (Fielding and Nicolson, 1980; Folk and Bradley, 2003).

Insects can slow or resist water loss through physiological means by inducing short-term changes to cuticle permeability (Bazinet et al., 2010), or by catabolising stored macromolecules (glycogen, lipid or protein) during desiccation, suggesting physiological responses to desiccation are energetically costly (Bosua et al., 2026; Tamang et al., 2022). In addition to yielding useful energy, catabolism of macromolecules, particularly of glycogen, is hypothesised to generate metabolic water for replenishing water lost during desiccation (Gefen and Ar, 2005; Halberg and Denholm, 2024). There is, however, limited supporting evidence for metabolic water production to be a primary driver or benefit of macromolecule catabolism in insects, but the contributions of stored macromolecules, and of nutrition in general, during and after desiccation show promising results for linking nutrition, metabolism and water loss to desiccation mortality (Archer et al., 2007; Bosua et al., 2026; Gefen and Ar, 2005; Gibbs et al., 1997; Halberg and Denholm, 2024).

A complete understanding of the physiological responses of insects to desiccation requires quantification of water loss and ion concentration or osmolality, ideally at the tissue-level scale to identify water compartmentalisation, but not all of these are reported in a study at this scale, or in conjunction with the restoration of homeostasis after recovery (Albers and Bradley, 2004; Machin, 1981). In addition, our mechanistic understanding of terrestrial arthropod water relations is limited because it primarily comes from a handful of specialist xeric arthropods (Insecta, Crustacea and Arachnida to name a few) that have provided examples of strong desiccation resistance evolved under extreme environments with limited opportunities for drinking water (Gefen and Ar, 2005; Naidu, 2001a). However, most arthropods are not xeric-specialists and studies on common models for insect physiology, including from selected lines of fruit flies (Archer et al., 2007; Folk and Bradley, 2003), have shown that although there are similarities in the response to desiccation employed by insects, such as in resisting or tolerating water loss, there is also high variation in these responses and in the underlying physiological mechanisms of desiccation resistance and tolerance within and among species and their habitat of origin (Addo-Bediako et al., 2001; Bosua et al., 2023; Hadley, 1994; Ramniwas and Kajla, 2012; Su et al., 2026). Overall, physiological processes occurring during desiccation can vary among taxa in where water is preferentially lost from body compartments, how ions are distributed between tissues, whether lost haemolymph is replaced with stored water from internal body compartments, as well as the extent to which excess haemolymph ions are sequestered or excreted (Gefen and Ar, 2005; Hyatt and Marshall, 1977; Koh and Wright, 2011; Nicolson et al., 1974; Wright et al., 1997).

In the case of the house cricket, *Acheta domesticus* (Orthoptera: Gryllidae), we know very little about its physiological response to desiccation and osmoregulation despite its history as a model insect species and use in mass-rearing industries (Woodring and Blakeney, 1980). Here, we tested the hypothesis that *A. domesticus* is a tolerator of water loss, whereby they preferentially lose water from the haemolymph over other body compartments during desiccation and tolerate water loss, in part, through active regulation of ion balance. If crickets actively resist water loss during desiccation, we predicted that observed water loss would be lower than expected from an exponential model representing passive water loss. We then tested a second hypothesis that this regulation is energetically costly, and fuelled by stored macromolecules (lipids, carbohydrates and proteins) during both desiccation and recovery. Over a time-course of desiccation and a bout of recovery, we measured water loss and water content of the whole body, gut, gonad, and carcass (largely muscle and cuticle). To measure changes in the haemolymph, we used ion-selective microelectrodes to determine whether crickets are regulating sodium or potassium ions in the hemocoel (Andersen et al., 2025), and used high molecular weight fluorescently-labelled dextrans to quantify haemolymph volume (Halberg and Denholm, 2024). To test whether desiccation and recovery is fuelled by stored macromolecules, we quantified whole-body total lipid, total protein and total glycogen. Together, these experiments point towards a two-phase strategy of crickets to tolerate a desiccation stress and be metabolically active during recovery. The results also reveal the relative importance of compartmentalised tissues for an insect’s water budget and desiccation resistance.

## Methods

### Experimental system

House crickets, *Acheta domesticus*, were obtained from a colony maintained at 30°C 14:10 h light:dark at Carleton University, ON, Canada since 2024. This colony was originally sourced as eggs from Aspire Food Group, ON, Canada. Colony crickets were housed in four plastic bins (68 L, 61 × 41 × 32 cm) of ∼200 individuals representing cohorts of approximately two weeks apart in age with egg cartons for shelter, and containers for feed and water. Crickets were fed coarsely ground chicken feed (layer pellets: 16 % crude protein, 2.5 % crude fat and 6.5 % crude fibre, Masterfeeds, ON, Canada). For each measured variable, adult crickets were sampled from the same colony bin until all uninjured individuals were removed, then the next oldest bin of adult crickets was sampled. Thus, where possible each experiment consisted of crickets from the same age cohort (1–2 months post adult ecdysis) and without visible physical injuries.

### Desiccation and recovery treatments

All crickets were sexed by the presence of an ovipositor and weighed (Mettler Toledo AB135-S, Ontario, Canada) at the start of the experiment to record their initial mass. We used five desiccation treatments: 0 (baseline), 3, 6, 18 hours of desiccation, and a recovery treatment (18 hours of desiccation then 6 hours of recovery with only water [starved]; 24 hours total). A total of four experiments were conducted using separate cohorts of individuals drawn from the colony: 1) ion concentration, 2) haemolymph volume, 3) combined tissue water and macromolecular content, and 4) tissue-specific water content. All experiments used the same desiccation treatments except for combined tissue water and macromolecular content, which had an additional recovery treatment (food and water [fed]). Crickets assigned to the 3, 6, 18 hour or either of the recovery treatments were placed in a 28 mL glass vial. The bottom of the vial had ∼10 mL Drierite (8 mesh indicating Drierite, W.A. Hammond Drierite Company, Ltd., OH, U.S.A.), which was separated from the cricket by cotton. The vial was capped with a washable sponge stopper. Vials (n = 10–16) were placed in a dark incubator (Incubator 4EM, Precision Scientific, IL, U.S.A.) for the desiccation treatment. A humidity probe placed inside a vial with a cricket and connected to a datalogger recorded humidity in the vial (5–15 % R.H.) and incubator temperature (31.2°C ± 0.04 °C s.e.) every 2 mins (IBS-TH2 Pro, Inkbird, Guangdong, China), giving a vapor pressure deficit of approximately 3.8 kPa.

At the end of the desiccation period, crickets were removed and weighed for their post-desiccation mass, which was used to calculate mass loss during desiccation (mg) and percentage change in initial mass (%). We assumed mass lost to be fluids but note this assumption overestimated water lost as some crickets were observed to excrete frass and sometimes oviposit. Crickets assigned to 0, 3, 6 or 18 hours desiccation were then processed to measure either their ion concentration, haemolymph volume, tissue-specific water or macromolecular content in one of the four experiments. Crickets assigned to either fed or starved recovery treatments were moved to an incubator (Isotemp refrigerated incubator, Thermo Fisher Scientific, ON, Canada, 30.06 °C ± 0.02 °C, 14:10 L:D, 86.7 % ± 0.4 % R.H.). These crickets were housed in a small container (96 mL, 7 cm diameter × 3 cm depth) containing a lidless 1.5 mL microcentrifuge tube capped with dental cotton for water, egg carton for shelter, and a feed dish made from a vial lid (1.7 cm diameter × 1.4 cm depth) glued on a 5 cm petri dish. If the cricket was assigned to the food and water recovery group, they were provided ground chicken feed. All crickets in the recovery group were removed after 6 h and weighed to record their post-recovery mass. Starved crickets had either their haemolymph ion concentration, haemolymph volume or combined tissue water and macromolecular content measured. Fed crickets had their combined tissue water and macromolecular content measured.

Crickets from all cohorts were pooled to characterise water loss during desiccation (273 females and 270 males). Dead crickets were dried in an oven for 48–72 hours (60 °C, Isotemp Oven senior model 203, Thermo Fisher Scientific, ON, Canada) then weighed to determine dry mass, either whole body or tissue component. Whole body dry mass of crickets dissected for tissue-specific water content was estimated from the sum of the dried guts, gonads and carcass. Whole body dry mass was subtracted from initial mass to calculate total body water content (mg). Water loss (%) was then calculated as the percentage of mass (assumed fluids) lost during desiccation relative to total body water content (Eqn 1). Water loss of 0 h desiccated crickets was 0 %.

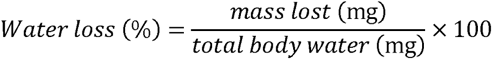

Eqn. 1. Water loss during desiccation calculated as a proportion of total body water lost during desiccation. Body water was calculated from oven dried whole body mass.

Mass loss was initially checked to see if a linear or exponential model best explained the data using Akaike Information Criterion (AIC) values. The exponential model fitted on natural log transformed data had lower AIC values than the linear model (5.66 fold difference in AIC values). Thus, subsequent analyses were conducted using natural log transformed on mass lost or water lost values.

### Combined tissue and tissue-specific water content

To partition body water between the haemolymph and tissues (gut, gonad and carcass including muscles combined; combined tissue water), crickets (n = 8 per treatment and sex) were anesthetised with CO_2_ for 10 seconds, then dissected by removing the head, hind legs and opening the abdominal and thoracic cavity. The internal tissues and leg muscles were blotted with delicate task wipes to absorb haemolymph. Crickets were then re-weighed to measure haemolymph-free wet mass. Gut contents were included but the presence of feed in the crop can bias mass and macromolecule estimations (Stabile et al., 2026). Crickets were then dried at 60 °C for 72 h and weighed to record their dried tissue mass. These dried individuals were then homogenised in a beadmill (Fisherbrand™ Bead Mill 24 Homogenizer, Thermo Fisher Scientific, ON, Canada) and stored at −80 °C until used for macromolecular analysis.

The amount of water (mg) in the whole body (pre-desiccation), tissues and the haemolymph (post-desiccation) was calculated by the difference between wet and dry mass of the respective body component. To control for among-individual differences in size, tissue water mass (mg) was converted into proportional body water content (%) following Eqn 2, interpreted as the amount of water remaining after desiccation.

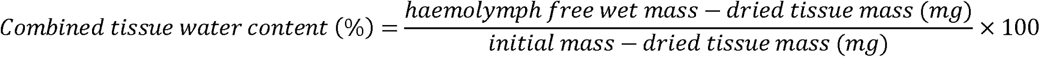

Eqn 2: Equation for calculating the proportion of body water in the tissue post-desiccation. The same equation is used for haemolymph substituting haemolymph free wet mass with whole animal wet mass.

To calculate the proportional amount of body water lost during desiccation between the haemolymph and tissues, the combined tissue or haemolymph water content (from Eqn 2) at 3, 6, 18, and recovery time points was subtracted from the mean values of the baseline group (0 h) for males and females separately. The 0 h group represents the baseline amount of body water partitioned between the tissue and haemolymph (0 % lost), where the difference represents the proportional amount of body water lost from baseline values with more negative values indicating greater water loss.

To partition combined tissue water further into tissue-specific water content, a second cohort of crickets (n = 8 per treatment and sex) were treated as described prior but instead the gut, gonads and remaining carcass were removed and weighed in separate pre-weighed 2 mL microtubes. The gut was ligated with cotton thread to preserve potential fluid in the gut lumen then gently lifted with forceps, excised and placed directly in the microtube without blotting. The gonads and carcass were blotted before weighing to obtain haemolymph-free wet mass. All tissues were oven dried then weighed for dry mass. Tissue-specific water content (%) was calculated as described above but using the mass of water in each specific tissue type (gonad, gut or carcass) and using the sum dry mass of all tissues as a proxy for dried tissue mass in Eqn 2. One male cricket appeared to have an impacted gut, resulting in an unusually heavy mass, and was removed from analysis.

### Haemolymph volume

To quantify haemolymph volume, we measured the dilution of a fluorescently labelled FITC-Dextran (FD4; 3–5 kDa, Sigma-Aldrich, St Louis, MO, USA), in the haemolymph using fluorescence spectrophotometry, modified from (Brzezinski and MacMillan, 2020). Crickets (n = 15–18 per treatment and sex) were anesthetised with CO_2_ then 1 µL of stock FITC-Dextran solution (4 mg mL^−1^) was injected into the ventral hind leg junction using a 10 µL Hamilton syringe. After 30 minutes, 10 µL of haemolymph was collected from each cricket by poking a 26G needle into the lateral prothorax-abdomen junction then squeezing gently to express a bead of haemolymph that was collected using a pipette, diluted x10 with anticoagulant buffer (44.8 mmol L^−1^ citric acid, pH 7.2), flash frozen in liquid nitrogen, and stored at −80 °C until use. Diluted haemolymph samples were pipetted into a 96-well plate and FITC-dextran content was quantified via fluorescence spectrophotometry (excitation: 485 nm, emission: 528 nm, BioTek Cytation 5 Imaging Reader, Winooski, VT, U.S.A). Haemolymph FITC-Dextran concentration was calculated from a standard range of 0.0008–0.2 mg mL^−1^ FITC-Dextran in anticoagulant buffer (x10 dilution), and haemolymph volume was calculated from the dilution of the FITC-Dextran stock solution.

### Haemolymph ion concentration

Haemolymph sodium (Na^+^) and potassium (K^+^) concentration at each time point was measured using ion-selective microelectrodes (Lajevardi et al., 2021). Thin-walled borosilicate glass capillaries (TW150-4, World Precision Instruments, Sarasota, FL, USA) were pulled to a tip diameter of ∼3–5 µm using a micropipette puller (P-1000, Sutter Instruments, Novato, CA, USA). K^+^-selective microelectrodes were backfilled with a 100 mmol L^−1^ KCl solution and K^+^ ionophore I cocktail B (Sigma-Aldrich). Na^+^-selective microelectrodes were backfilled with a 100 mmol L^−1^ NaCl solution and Na^+^ ionophore X (Sigma-Aldrich). The reference electrode was a pulled filamented glass capillary (1B200F-4, World Precision Instruments, Sarasota, FL, USA) filled with 500 mmol L^−1^ KCl. Microelectrodes were calibrated using 3 and 30 mmol L^−1^ KCl for K^+^ electrodes and 30 and 300 mmol L^−1^ NaCl for Na^+^ electrodes, with osmolality adjusted with LiCl.

Haemolymph was collected (n = 17–28 per treatment and sex for each ion) as described previously and placed under hydrated paraffin oil in in a Petri dish lined with silicone elastomer (Sylgaard 184 Silicone Elastomer Kit, Dow Chemical, Midland, MI, USA). Ion concentration was calculated from the measured voltage using the equation, *C_f_* = *C_L_*10^*ΔV/S*^ where *C_f_* is the final concentration (mmol L^−1^), *C_L_* is the concentration (mmol L^−1^) of the lowest standard for the respective ion, *ΔV* is the difference in voltage (mV) between the sample and the lowest standard, *S* is the difference in voltage (slope, mV) between the two standards. Microelectrodes with a slope close to the Nernst slope of 58 mV were used (50- –66 mV, mean electrode slopes: K^+^ = 54.4 ± 0.08 mV, Na^+^ = 57.6 ± 0.1 mV).

### Macromolecule use

#### Total lipid content

Total lipid content was quantified using the vanillin assay adapted from (Haider et al., 2018). Aliquots of dried ground tissue (5–15 mg) from each cricket was mixed with 1 mL chloroform-methanol 2:1 v/v and agitated for 30 min on a plate rocker. Samples were centrifuged (3000 × *g*, 5 min) with 210 µL 0.25 % KCL (w:v) to induce phase separation. A 150 µL aliquot of the organic layer was evaporated at 95 °C before 60 µL of concentrated sulphuric acid was added and the capped tube was incubated at 95 °C for 10 mins. The samples were cooled to 21 °C and 1.4 mL vanillin reagent (1.2 % vanillin w/v in 85 % phosphoric acid) was added and left to incubate at room temperature for 5 min. Aliquots of 200 µL were added to a 96-well microplate in triplicate and absorbance at 490 nm was read using a microplate reader (Epoch, BioTek, Winooski, VT, U.S.A.). Sample concentrations were calculated from a standard curve of corn oil in acetone (0.2–4.5 mg mL^−1^). Lipid content was calculated as mg lipids in whole dried tissue mass.

#### Protein content

Protein content was measured using the bicinchoninic acid (BCA) assay (Pierce BCA Protein Assay Kit, Thermo Fisher Scientific, ON, Canada). Dried tissue aliquots of 3–5 mg were homogenized in 1 mL ice-cold buffer (0.1 mol L^−1^ Tris-HCl pH 8.0, 0.2 % Triton X-100, 153 µmol L^−1^ MgSO_4_ and 0.1 mmol L^−1^ PMSF) in a beadmill. The homogenate was centrifuged (3000 × *g*, 10 min, 4 °C) then a 100 µL aliquot of the spun homogenate was mixed with 0.8 mL working reagent (1:8) and incubated at 37 °C for 30 min. Aliquots of 225 µL of the mixed solution was added to a 96-well microplate in triplicate, then read at 562 nm using a microplate reader. A standard curve (0.125 – 2.0 mg mL^−1^) was generated using bovine serum albumin (Sigma-Aldrich, ON, Canada) diluted in the complete homogenizing buffer. Protein content was calculated as mg protein in whole dried tissue mass.

#### Glycogen content

Glycogen content as a proxy for carbohydrates was measured using the anthrone assay, adapted from (Toxopeus et al., 2016). Dried homogenised tissue aliquots of 15–18 mg were heated in 1 mL KOH (30% KOH w/v) at 90 °C for 30 min. The homogenate was centrifuged (3000 × *g*, 10 min, 21 °C) then 500 µL of the supernatant was mixed with 25 µL of saturated Na_2_SO_4_ and 1 mL of ethanol (95%). This solution was centrifuged again then the supernatant discarded and the precipitated pellet desiccated at 90 °C for 30 min. The dried pellet was resuspended in 250 µL dH_2_O then mixed with 750 µL of anthrone reagent (0.5 g L^−1^ anthrone in 72% sulphuric acid) and incubated at 90 °C for 15 min. The cooled solution was added to a 96-well microplate in triplicate, then read at 620 nm using a microplate reader. A standard curve (0.1 – 1.0 mg mL^−1^) was generated using oyster glycogen II diluted in dH_2_O. Glycogen content was calculated as mg glycogen in whole dried tissue mass.

### Data analysis

Data was analysed in R v4.5. All data is presented as means ± standard error. Ion concentration was Log_10_ transformed prior to analysis to better meet the assumptions of linear regression. Linear regressions used desiccation treatment (0, 3, 6, 18, 24 h starved & 24 h fed) and sex as fixed categorical predictors except for mass lost and water lost, which used continuous variables. Initial models for regressions, except for macromolecules, included the interaction between desiccation treatment and sex, which was dropped from further analysis if the interaction was not significant. No interaction between time and sex was found for mass lost, water loss rates, potassium concentration, and tissue-specific water content. An interaction between time and sex was present for sodium concentration and haemolymph volume. To account for cricket size, initial mass was used as a covariate of haemolymph volume. An initial model for combined tissue water content showed sex and an interaction between sex and time was not significant, thus sexes were pooled. Diagnostic plots checked the model assumptions were met. Total lipid or carbohydrate content was a mixed effects model starting with a full model including starting mass, time as a categorial variable, sex, and their interactions, as well as plate identity as a random variable. Models were fitted using Maximum Likelihood using *lme4* and backwards stepwise model selection conducted based on AIC values using *lmerTest* to determine the reduced model (Supplementary Material). Total dry mass and protein content were analysed using a fixed model because the random plate effect was not significant. Initial wet mass was used as the covariate to account for variation in mass between sampled cohorts assigned to time treatments (Fig. S2). The three-way interaction between initial mass, time and sex was not significant in all models. Post-hoc pairwise comparisons were made using Tukey’s HSD on estimated marginal means (*emmeans* package) for the within-sex effect if there was an interaction or on the pooled effects of sex if there was no interaction. There were strong and consistent differences between males and females in all measured responses except for total lipid, glycogen content and combined tissue water content.

## Results

### Total body water content and loss during desiccation

Both models for wet mass lost and water loss during desiccation underestimated mean values at each time point in males and females (Fig. 1). The random effect of experimental cohort had a variance of 0.002 ± 0.044 s.d. for the model for mass lost and of 0.002 ± 0.048 s.d. for the model for water loss. Mean mass loss (% of initial mass) at 18 h was 9.941 % ± 0.347 for females and 16.158 % ± 0.0.368 for males. Crickets lost mass throughout desiccation (Fig.1A, GLMM, t_542.6_ = 36.210, P < 0.001) and males consistently lost more mass than females (Fig. 1A, t_539.8_ = 16.208, P < 0.001). Mean total water loss (% of total body water) at 18 h was 14.517 % ± 0.452 for females and 21.101 % ± 0.460 for males. Water loss increased over time (Fig. 1B, GLMM, (GLMM, t_542.2_ = 37.885, P < 0.001) and water loss was higher for males than females (t_539.4_ = 13.672, P < 0.001). Crickets regained mass during recovery but not to pre-desiccation levels (Supplementary material, Fig. S1).

**Figure 1.**
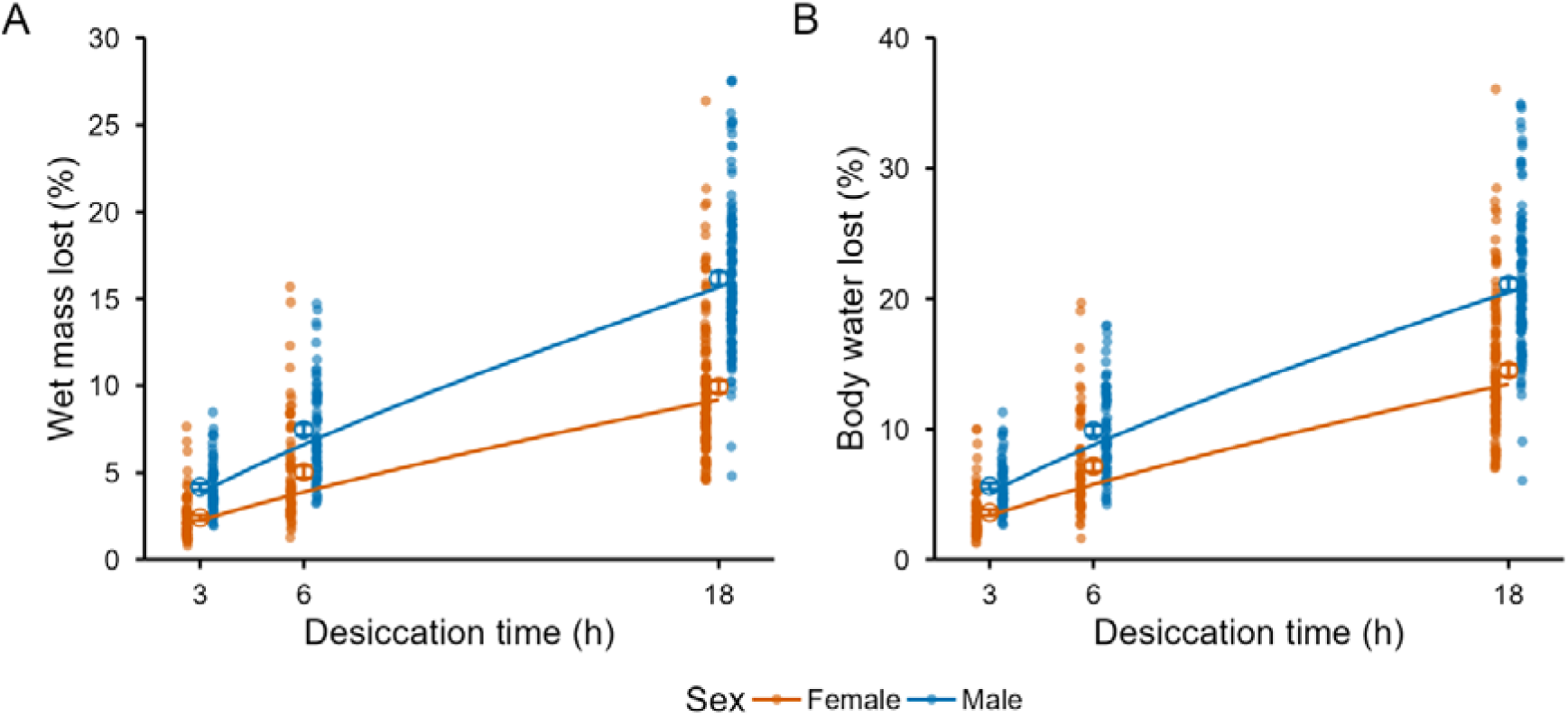
Crickets (*Acheta domesticus*) progressively lose water during desiccation stress. A) Percentage of initial mass lost after desiccation (h). B) Water loss (% total body water) during desiccation. Orange points are females (n = 273). Blue points are males (n = 270). Open circles with error bars represent mean ± sem. Lines are best fitted exponential models on the fixed effects representing expected non-linear mass or water loss relationships. Recovery groups were pooled with 18 h groups using post-desiccation mass.

### Tissue water content

Crickets in the combined tissue water content cohort, pooling sex, lost up to 15.417 ± 1.228 % of their body water over 18 h of desiccation (Fig. 2A). This loss was almost equally split between loss from the tissues (guts, gonads and carcass combined) and from the haemolymph (53 and 47 % from the tissues at 3 and 18 h) but was only 11 % from the tissues at 6 h. Post-hoc tests indicated after recovery, crickets restored their haemolymph water content to baseline levels but were only able to restore tissue hydration state if provided both food and water (Fig. 2A, starved recovery group: t_90_ = 3.728, P = 0.004). Water loss from specific tissues over time were similar between sexes and was mostly significantly different after prolonged desiccation (Fig. 2B). Gut water content decreased after 18 h and after recovery (ANOVA, F_(4, 74)_ = 8.7, P < 0.001), gonad water content did not change over time (ANOVA, F_(4, 74)_ = 1.5, P = 0.212), and carcass water content decreased after 18 h and after recovery (ANOVA, F_(4, 74)_ = 7.8, P < 0.001).

**Figure 2.**
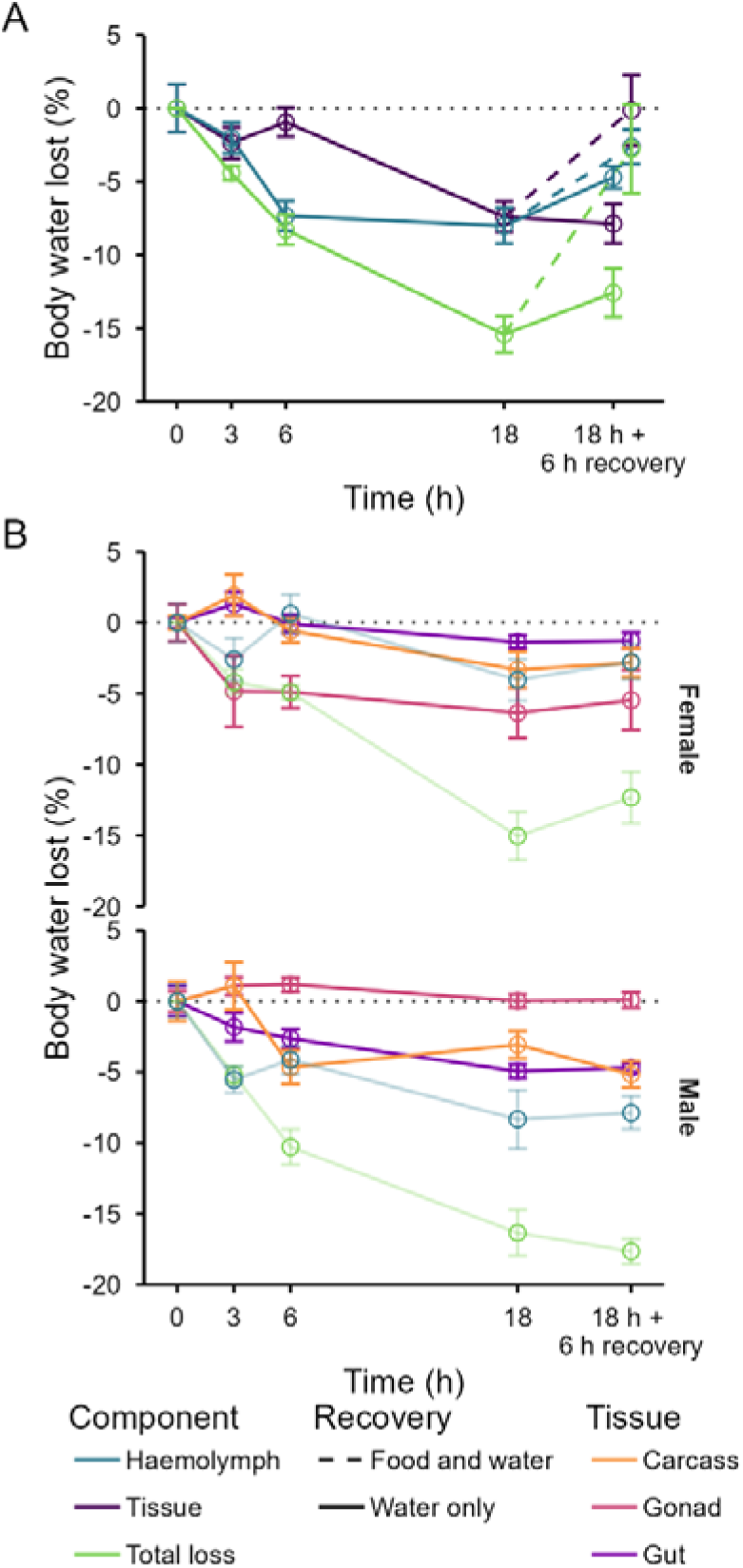
Relative water content across different body components decreases throughout a desiccation event and after a bout of recovery relative to the mean of the control (0 h) crickets. Relative body water content (%) was calculated relative to the pre-desiccation water content of an individual. A) Whole body water loss (green) is separated into water loss from the tissues (purple; gut, gonad and carcass combined) and from the haemolymph (blue). Males and females are pooled (n = 16 per time point). Solid lines are starved crickets and those provided only water during recovery. Long dashed lines are crickets provided food and water during recovery (24 h). B) Water loss from the tissues partitioned into tissue-specific components (n = 8 per time point and sex, second cohort of *A. domesticus*). Open circles with error bars represent mean ± s.e..

### Haemolymph volume

Changes in haemolymph volume over time measured using FITC-Dextran dilutions varied between sexes (Fig. 3A, ANOVA, F_(4, 153)_ = 6.444, P < 0.001), where post-hoc tests indicated haemolymph volumes of males did not change throughout. Haemolymph volume in females did not differ between 0 h and 6 h of desiccation, then decreased after 18 h compared to baseline levels (t_153_ = 6.484, P < 0.001). Haemolymph volume of females then increased after recovery compared to 18 h (t_153_ = −3.830, P = 0.002) and returned to baseline levels (t_153_ = 2.486, P = 0.099).

**Figure 3.**
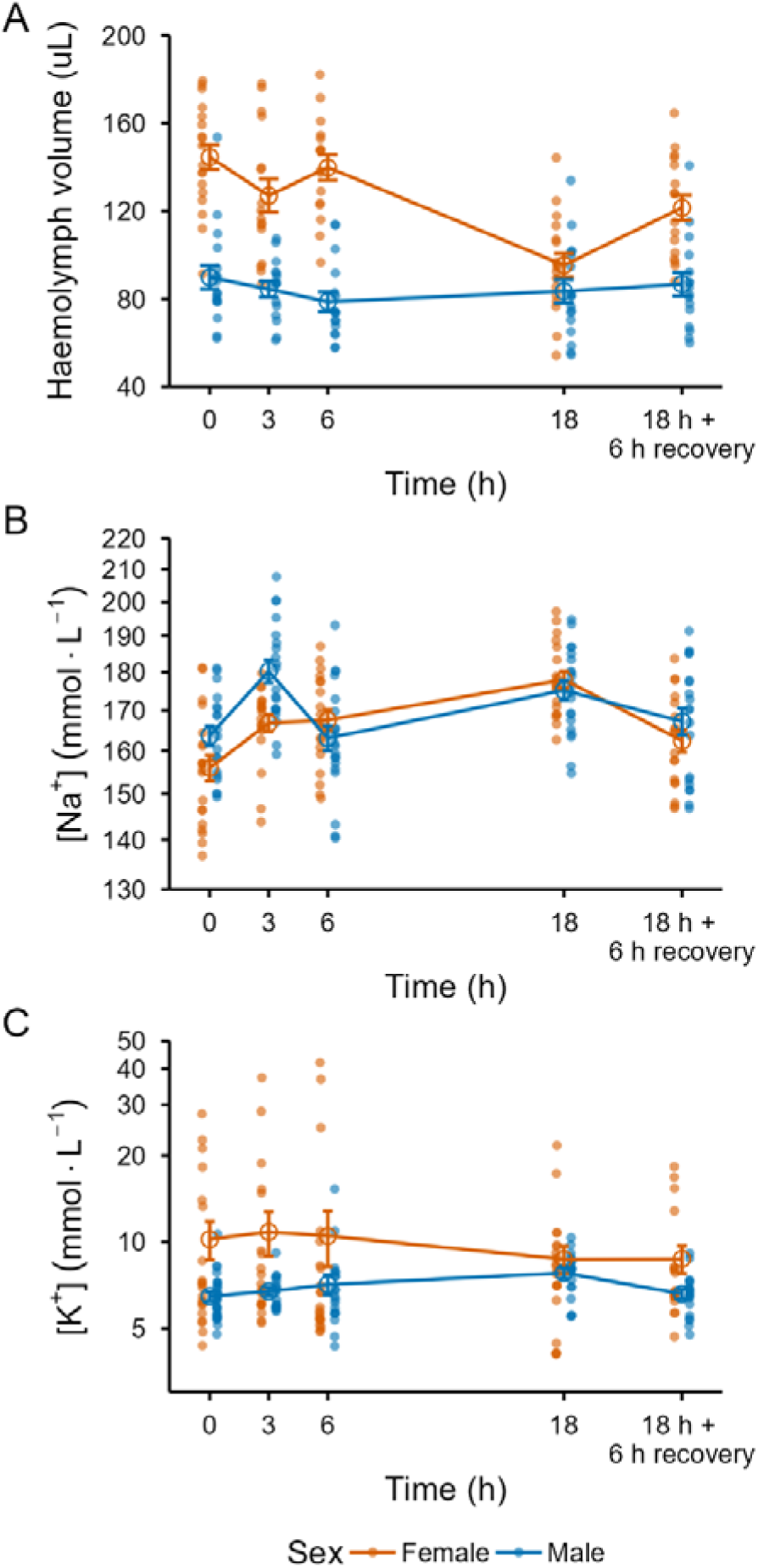
A) Haemolymph volume (uL) of desiccated crickets (n = 15–18 per sex and time). B) Sodium (Na^+^) and C) potassium (K^+^) ion concentration (mmol·L^−1^) for desiccating crickets (n = 17–28 per sex and desiccation time). Orange points are females. Blue points are males. Points with error bars are standard error about the mean connected by lines.

### Haemolymph ion concentration

Changes in sodium concentration over time varied between sexes (Fig. 3B, ANOVA, F_(4, 183)_ = 3.636, P = 0.007). Sodium concentration increased over 3h in males and females then decreased in males after 6 hours to baseline concentrations (HSD, t_183_ = 0.221, P = 1) but continued to increase in females (Fig. 3B). Sodium concentrations were elevated above baseline at 18 h of desiccation for both females (HSD, t_183_ = −5.802, P < 0.001) and males (HSD, t_183_ = −2.929, P = 0.031) and returned to baseline levels after rehydration in females (HSD, t_183_ = −1.779, P = 0.389) and males (HSD, t_183_ = −0.828, P = 0.922). Baseline haemolymph potassium concentrations were higher in females than in males (Fig. 3C, ANOVA, F_(1, 195)_ = 12.742, P < 0.001) but did not change during desiccation or recovery (F_(4, 195)_ = 0.399, P = 0.809). Five crickets were unable to be measured because they did not have sufficient volumes of haemolymph. Two of these individuals (one was sampled after 3 h and the other after 18 h desiccation) were active after desiccation and displayed an escape response. The other three individuals all experienced 18 h of desiccation and were lethargic. All were unable to recover when given water and eventually died.

### Macromolecule content

There was an interaction between time and sex in the reduced model of dried tissue mass (guts, gonads and carcass combined, Fig. 4A and Fig.S2, F_(5, 82)_ = 8.711, P < 0.001). This trend was driven by starved female crickets (HSD, P < 0.05), whose mean dry mass decreased to 151.654 ± 10.801 mg compared to 213.536 ± 19.191 mg in control females, and by fed male recovery crickets that were heavier than starved male recovery crickets (t_82_ = 3.031, P < 0.001). In the reduced linear models for each macromolecule there was an overall effect of time on total lipid, total protein (F_(5, 82)_ = 6.987, P < 0.001), and total glycogen (F_(5, 89.8)_ = 5.574, P < 0.001). An interaction between time and sex was only present in the reduced lipid model (F_(5, 91.9)_ = 2.525, P = 0.035). These relationships were driven by differences between the recovery group and other desiccation time points. Total lipid content of starved female crickets after recovery were lower than 18 h female crickets (Fig. 4B, t_119.4_ = 3.146, P = 0.025). Starved recovery crickets had lower protein content than crickets after 3 hours of desiccation (Fig. 4C, t_82_ = 3.391, P = 0.013) and 18 hours of desiccation (t_82_ = 3.442, P = 0.011). Fed recovery crickets had higher glycogen contents than 6 h desiccated crickets (Fig. 4D, t_100.2_ = −3.083, P = 0.031) and starved recovery crickets had lower glycogen contents than the 3 h group (t_99.2_ = 3.272, P = 0.018) and the fed recovery group (t_99.3_ = 4.335, P < 0.001).

**Figure 4.**
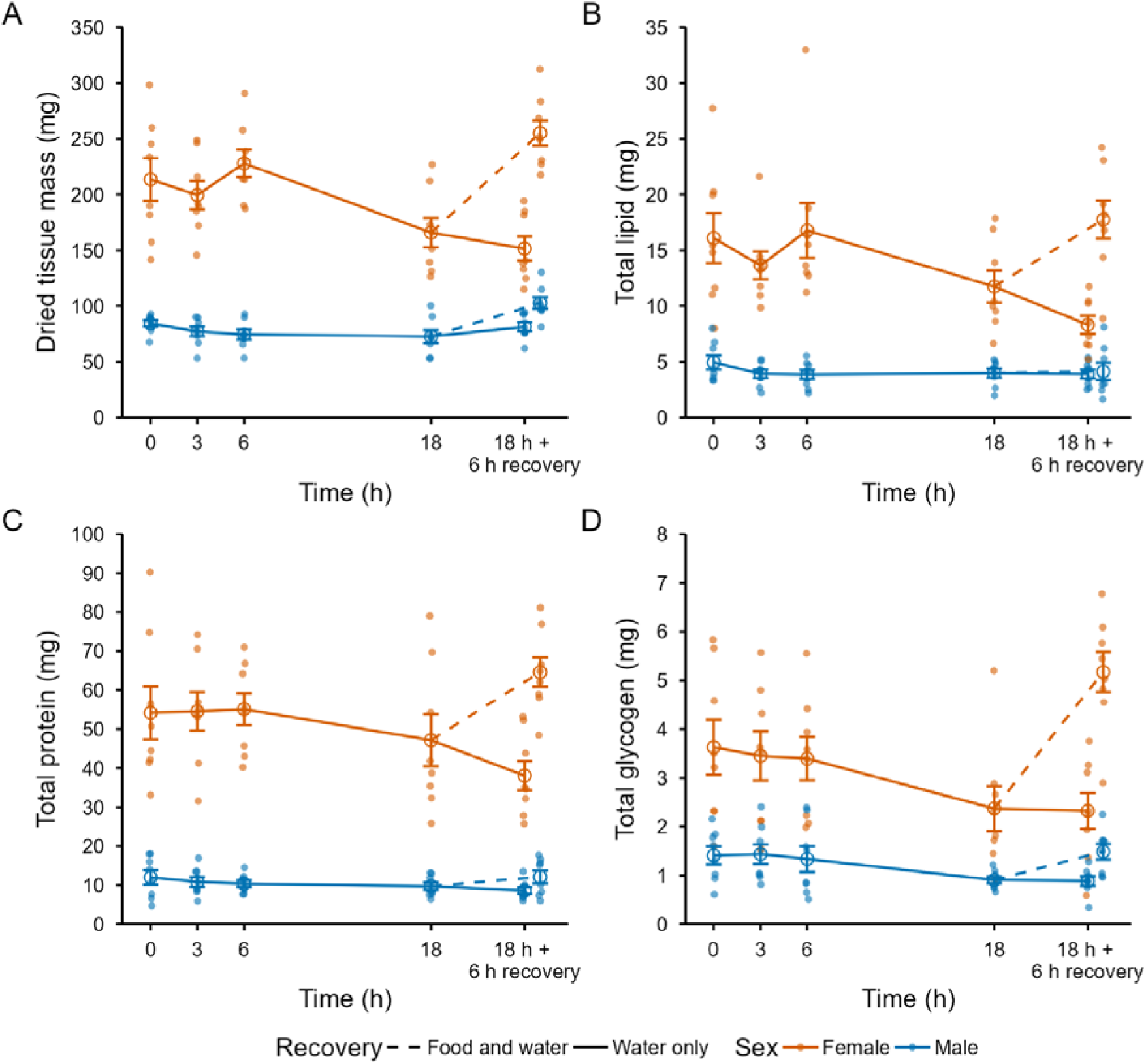
Macromolecular use is limited throughout the time-course of desiccation and used during recovery. A) Dried tissue mass (mg), B) Total lipid content (mg), C) Protein content (mg), and D) glycogen content (mg) of desiccated female (orange points, n = 8 per time point) and male (blue points, n = 8 per time point) crickets. Points with error bars are standard error about the mean connected with lines. Solid lines are starved crickets and those provided only water during recovery. Dashed lines are crickets provided food and water during recovery (18 h desiccation and 6 h recovery).

## Discussion

To understand the response and recovery of insects to desiccation, we have quantified the time course of key ionoregulatory and osmoregulatory mechanisms during desiccation and recovery in a commonly studied mesic insect species. We conclude that house crickets *A. domesticus* are tolerators of water loss, employing different strategies to manage water loss between sexes, and have distinct physiological mechanisms occurring during and after desiccation. House crickets notably differ from other species in three ways: 1) water loss is non-discriminatory between the haemolymph and the tissues during prolonged desiccation and crickets do not actively resist water loss, 2) crickets are ionoregulating during desiccation but this is not fuelled by stored macromolecules, and 3) recovery near complete and energetically costly in its use of stored macromolecules. Studies in arid adapted and mesic insects to date have shown there are broad generalities in the response phenotype across taxa in how individuals can resist or tolerate water loss, as well as maintain homeostasis during a stress event, but also a large diversity in how these phenotypes are achieved through the relative contribution of various physiological mechanisms underlying the desiccation response (Albers and Bradley, 2004; Johnson et al., 2026; Koh and Wright, 2011; Wright et al., 1997). Thus, there is diversity in how insects can be tolerators or resistors of desiccation occurring over different time scales that are taxon and sex specific yet achieves a similar phenotype.

The time course of desiccation responses is notable because desiccation events may last from minutes to hours to days and this is expected to change with climate change. An individual’s capacity to withstand future desiccation stress may depend on their tolerance or resistance to desiccation which, in turn, is determined by a species’ evolutionary history or ecology. For example, it takes 6 days for a desert dwelling beetle to lose the same proportion of initial mass that a cricket here loses in 18 hours under similar temperatures and relative humidities (Naidu, 2001a). This diversity becomes important when inferring potential responses to climate change and relative vulnerability to desiccation stress as these time-dependent differences can result in vastly different consequences for individuals of different species.

### Crickets tolerate desiccation

Water loss rate was best explained by a negative exponential relationship that suggests evaporative water loss proceeds slightly more slowly as the osmolality of the haemolymph increases (Fig. 1). In fact, the model underestimated water loss rates at 6 hours of desiccation, suggesting crickets do not appear to be continuously resisting water loss throughout this period even if an initial resistance occurred. Water loss could be resisted by closing spiracles or reducing ventilation in ways we have not measured. This can be confirmed by using high-resolution flow through respirometry, to partition out respiratory and cuticular water loss (Bosua et al., 2023). Males had more body water proportional to mass, lower haemolymph volumes and higher rates of water loss than females. Both sexes had a similar proportion of haemolymph volume or mass accounting for whole body wet mass. These results suggest that it is easier for males to lose water than females or that they do not prevent water loss as much as females, perhaps because the eggs in the haemocoel of females reduces the effective volume of the haemocoel and the volume of freely circulating haemolymph that can be readily lost. Sex specific differences in cuticle permeability could explain the differences in water loss between sexes, as reported in alfalfa cutting bees where higher male cuticle permeability was associated with higher water loss rates than females (Johnson et al., 2026).

### Water loss among body compartments changes over the course of desiccation

An unexpected result was that water loss was not consistently partitioned among body compartments throughout desiccation. Although water loss was predominantly from the haemolymph at 6 h, by 18 h the cumulative loss was approximately equally divided between haemolymph and tissues when sexes were pooled (Fig. 2A). Most previous studies have concluded that the haemolymph is sacrificed in favour of maintaining intracellular water volume and ion concentration in insects (Nicolson et al., 1974; Wright et al., 1997). This result becomes more complex when comparing with our FITC-Dextran results which show that females have reduced haemolymph volume after 18 h while males do not (Fig. 3A). The difference could reflect the method used as haemolymph volume measured using FITC-Dextran may include parts of the haemocoel that are harder to access through dissection, whereas haemolymph water loss measured gravimetrically is proximately calculated from the subtraction from total body water. At a tissue-specific level, our gravimetric estimates suggest males are losing approximately two thirds of their lost body water during desiccation from the haemolymph compared to their carcass, gut or gonad, whereas females are losing a more equal mix from these body compartments at 18 h (Fig. 2A vs Fig. 3A).

Arthropods can compartmentalise water loss which likely serves an adaptive purpose for life on land (Machin, 1981). At a tissue-specific level, the largest changes in water loss occurred from the gut. The loss could be occurring from water in the gut lumen, which then suggests that most water loss in crickets occurs from extracellular spaces (haemolymph and gut lumen). We observed that some crickets had crops full of liquid during dissection, usually after recovery and likely from drinking. If cricket crops are permeable to water, then the use of the crop to store water and replenish lost haemolymph is possible for hydrated individuals (Serrao et al., 2014). The decrease in gut water was not strongly associated with changes in haemolymph volume (comparing Fig. 2B with Fig. 3A) in females as a decrease in gut water content did not correspond with maintenance of haemolymph volume over time. However, in males the decreasing gut water content while maintaining haemolymph volume could indicate that water in the gut lumen is preferentially lost during desiccation. This process has been described in scorpions that use stored water in their hepatopancreas during desiccation, where haemolymph volume will only decrease after the hepatopancreas is depleted (Gefen and Ar, 2005). Alternatively, this water could be excreted from the gut. The lack of water loss from the muscles and carcass and gonads suggests these intracellular spaces are protected during prolonged desiccation. This protection is notable because replacing lost tissue water appears to be slower than replenishing lost haemolymph water during recovery (Fig. 2A). Female eggs and ovaries had low water contents (mean 48.75 % ± 0.3 s.e.) and the desiccation stress was likely not chronic enough for egg reabsorption to be beneficial.

### Crickets are strong ionoreguators during desiccation and recovery

House crickets ionoregulate despite considerable loss of body water during desiccation. Like in many other arthropods, loss of haemolymph volume did not result in a proportional increase in haemolymph ions in crickets with an expected regression slope of 1 between changes in sodium concentration and changes in body water from baseline (Folk and Bradley, 2003; Naidu, 2001b). Rather, the slope of this relationship was 0.022 on log-log axes, pooling sex, suggesting that crickets are ionoregulating during desiccation in the face of water loss (Fig. S2). Water loss and a reduction in haemolymph volume did not induce hyperkalaemia and crickets were able to regulate their potassium ion concentrations despite a reduction in haemolymph volume, which is also found in other insects (Albaghdadi, 1987; Hyatt and Marshall, 1985; Naidu, 2001a). Although some females had unusually high potassium concentration, this was not correlated with desiccation duration, cohort or the amount of water loss, and could be from injury to tissues or eggs during haemolymph extraction. Overall, males appeared to regulate sodium more strongly than females which showed consistent increases in haemolymph sodium, despite males experiencing more water loss. If there was excess sodium in the haemolymph, then what happened to it? There are three possible fates to this sodium. Excess sodium could be exchanged between the intracellular and extracellular spaces or removed by excretion or sequestration into salts (Folk and Bradley, 2003; Hyatt and Marshall, 1977; Naidu, 2001a; Wright et al., 1997). We observed a spike in sodium concentration in males at 3 h, which returns to baseline concentrations at 6h. This result is unlikely to be experimental variation because it was repeated between two independently measured cohorts and we suggest this spike could originate from sodium in the tissues, which is transported into the haemolymph to maintain intracellular osmolality or be excreted. The exact fate of ions is sometimes unclear in other studies (Koh and Wright, 2011). The development of tools for studying osmoregulatory mechanisms can provide opportunities to quantify the movement of water and ions within the body during desiccation at resolutions previously unavailable (Andersen et al., 2025). This work can be expanded with direct measurements of osmolality and tissue-specific ion concentration including of the gut lumen contents, as well as the function of the renal system under acute stress.

### The desiccation response is a two-stage period of inactivity and a costly recovery

Our results reveal that recovery is not simply the reversal of processes occurring during desiccation but involves its own mechanisms and associated energetic costs indicated by reductions in dried tissue mass and macromolecules (Fig. 4). Recovery was also asynchronous among physiological compartments. Haemolymph volume and ion concentrations returned to baseline during the 6-h recovery period, whereas restoration of tissue hydration depended on access to food as well as water. Thus, recovery of haemolymph and ion homeostasis can be decoupled from restoration of tissue water content. This can be interpreted as recovery itself is costly or that the desiccation response is costly, but such costs are delayed being paid for during recovery. These interpretations are not mutually exclusive, and a compounding of desiccation responses with recovery costs would explain the consist differences in stored macromolecules between starved recovery crickets and desiccated crickets. The lack of a corresponding decrease in dried tissue mass and stored macromolecules of fed recovery crickets suggests that feeding is replenishing stored macromolecules and masking the apparent cost of desiccation. Ion concentrations, haemolymph volume and tissue dry mass were all similar to baseline levels after 6 h of recovery, suggesting the magnitude of the stress was not excessive to cause lasting detrimental effects for an individual or for an individual to fail to restore homeostasis, even if crickets did not fully regain their initial wet mass. Our macromolecular results suggest that this restoration of ion homeostasis is fuelled by stored macromolecules. Similar capacities for recovery are widely reported in other insects but have not reported on the explicit role of food provision during recovery (Albers and Bradley, 2004; Benoit et al., 2005; Folk and Bradley, 2003; Hyatt and Marshall, 1977; Hyatt and Marshall, 1985; Naidu, 2001a; Naidu, 2001b).

We found no support for the production of metabolically-derived water during desiccation, particularly from glycogen (Archer et al., 2007; Djawdan et al., 1998; Gibbs et al., 1997). In fact, crickets did not expend significant stored macromolecules during desiccation. This suggests osmo- and ionoregulation during desiccation is not metabolically intensive and crickets are likely inactive while confined in the glass vials. Some individuals were more active during desiccation than others as some crickets had chewed into the cotton or sponge in the vial. Lipid and protein content trended downwards in at 18 h in females but this trend was not significant (Fig. 4). This trend was likely driven by a cohort effect because these individuals were smaller at the start. Changes in glycogen during desiccation have been suggested to indicate that metabolically stored water is liberated in direct response to the desiccation (Bosua et al., 2026; Djawdan et al., 1998) (but see (Hoffmann et al., 2001)). Stored proteins and lipids have also been shown to be used during desiccation in some insects, where the relative use of carbohydrates, proteins or lipids differ between during an acute stress and during recovery (Archer et al., 2007; Gefen and Ar, 2005; Naidu, 2001a).

### Conclusions

We have established a baseline for how a commonly studied mesic species can withstand up to 18 hours of a severe desiccation, and with minimal physiological consequences in the variables measured here for up to six hours of desiccation. This information suggests crickets could be resilient to a potential desiccation event due to extreme weather provided the stress is shorter than 18 hours and they can adequately recover in six hours. In nature, insects can respond to desiccation through morphological or behavioural means, such as by closing their spiracles to reduce evaporative water loss at the expense of gas exchange or to retreat to humid microclimates (Bosua et al., 2023; Pincebourde and Woods, 2020). As house crickets are an important species in the edible insect industry, our results could be important for designing how water is provided in mass-rearing methods as this is a challenging balance between not providing enough water, resulting in desiccation stress, and providing too much, increasing the risk of drowning, fungal and bacterial growth and the spread of disease (Kong et al., 2026; Morales-Ramos et al., 2024).

## Data accessibility statement

The datasets and code supporting this article have been uploaded to Zenodo doi: 10.5281/zenodo.23360579.

## Author contributions statement

JDK: Conceptualization, Data curation, Formal analysis, Investigation, Visualization, Writing – original draft, Writing – review & editing. SMB & HAM: Funding acquisition, Methodology, Writing – review & editing

