## Supplementary Material for "Energetic costs of osmo- and ionoregulatory responses to desiccation and recovery in a model insect"

### Post-desiccation recovery

Table S1. Mean (± standard error) change in wet mass at the start and end of a desiccation stress (18 h) and after recovery (6 h). Crickets fed or starved during recovery are pooled.

| Sex | n | Pre-desiccation | Post-desiccation | Post-recovery |
| --- | --- | --- | --- | --- |
| Female | 58 | 100 | 91.229 ± 0.434 | 94.357 ± 0.635 |
| Male | 60 | 100 | 82.876 ± 0.588 | 88.604 ± 0.966 |

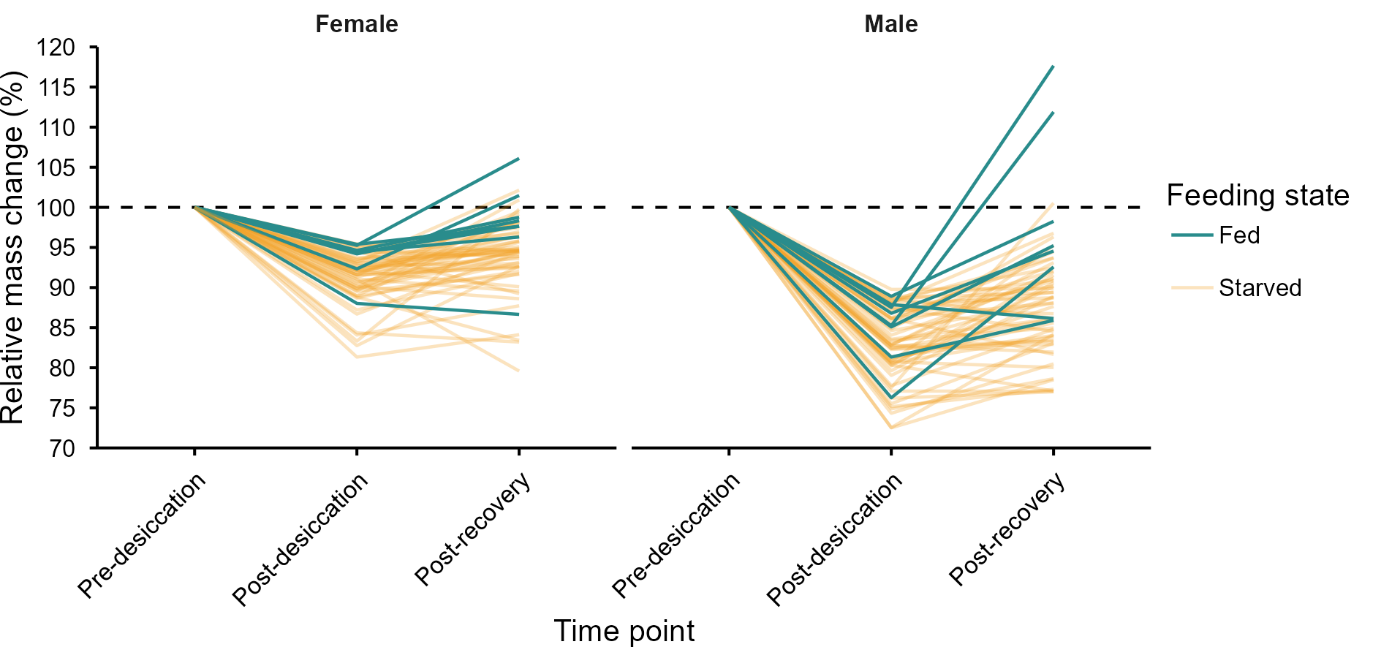

Figure S1. Crickets (n = 58 females and 60 males) lose mass during 18 h of desiccation and most individuals gain mass after 6 h of recovery with water and with (green lines) or without (gold lines) food. Lines are individual crickets. Mass change was calculated relative to initial mass (pre-desiccation).

### Relationship between ion concentration and body water

We tested the hypothesis that hemolymph ion concentration increases proportionally to body water lost during desiccation. As water loss is expected to follow an exponential relationship, we natural log transformed body water loss and ion concentration (K^+^ or Na^+^) to fit a linear regression for 3, 6, and 18 h desiccated crickets (n = 116 for potassium and 117 for sodium). We fit an initial full model with sex and the interaction between sex and water loss for both ions. Sex was not significant in both models and the term was removed from the final model.

Potassium ion concentration was independent of body water loss (Fig. S2A), rejecting the hypothesis, with an equation of Log[K^+^] = Log Water loss × −0.016 + 2.068 (slope s.e. ± 0.052, t_114_ = −0.308, P = 0.759, R^2^ = 0.08 %). Sodium ion concentration increased with increasing water loss (Fig. S2B) following the equation Log[Na^+^] = Log Water loss × 0.022 + 5.092 (slope s.e. ± 0.010, R^2^ = 4.024 %).

The slope for Na^+^ was different to 0 (t_115_ = 2.196, P = 0.030). We further tested if this slope differed from 1 by fitting a linear model in the form of (Log[ion] – Log Water loss) = Log Water loss. The offset model had a slope different to 1 (slope = −0.977, t_115_ = -−93.900, P <0.001, R^2^ = 98.71%).

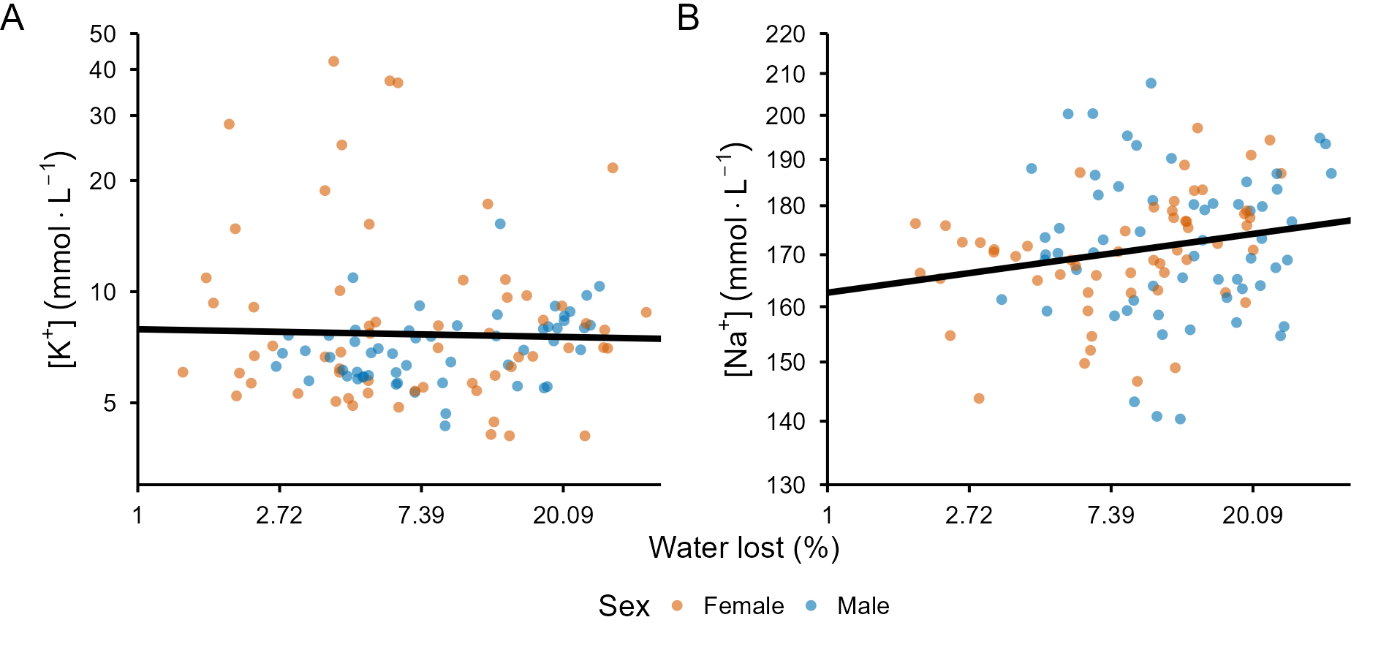

Figure S2. A) Haemolymph potassium (K^+^, n = 116) does not change with body water lost but B) haemolymph sodium (Na^+^, n = 117) increases with water lost with a slope of 0.022. Male (blue) and female (orange) crickets desiccated for 3, 6, and 18 hours were pooled for analyses. Black line represents the fitted linear regression. Axes are natural log transformed.

### Stored macromolecules analyses

To check the relationship between initial mass at the start of the experiment and dried tissue mass, we fitted a linear model with sex and the interaction between sex and initial mass as covariates (Fig. S3). The interaction term was significant (t_92_ = -2.880, P = 0.005), giving the equations: Female dried tissue mass = initial mass × 0.382 - 34.885 and male dried tissue mass = initial mass × 0.253 - 4.345. Coefficient of determination (R^2^) was 95.4 % for 48 males and 48 females.

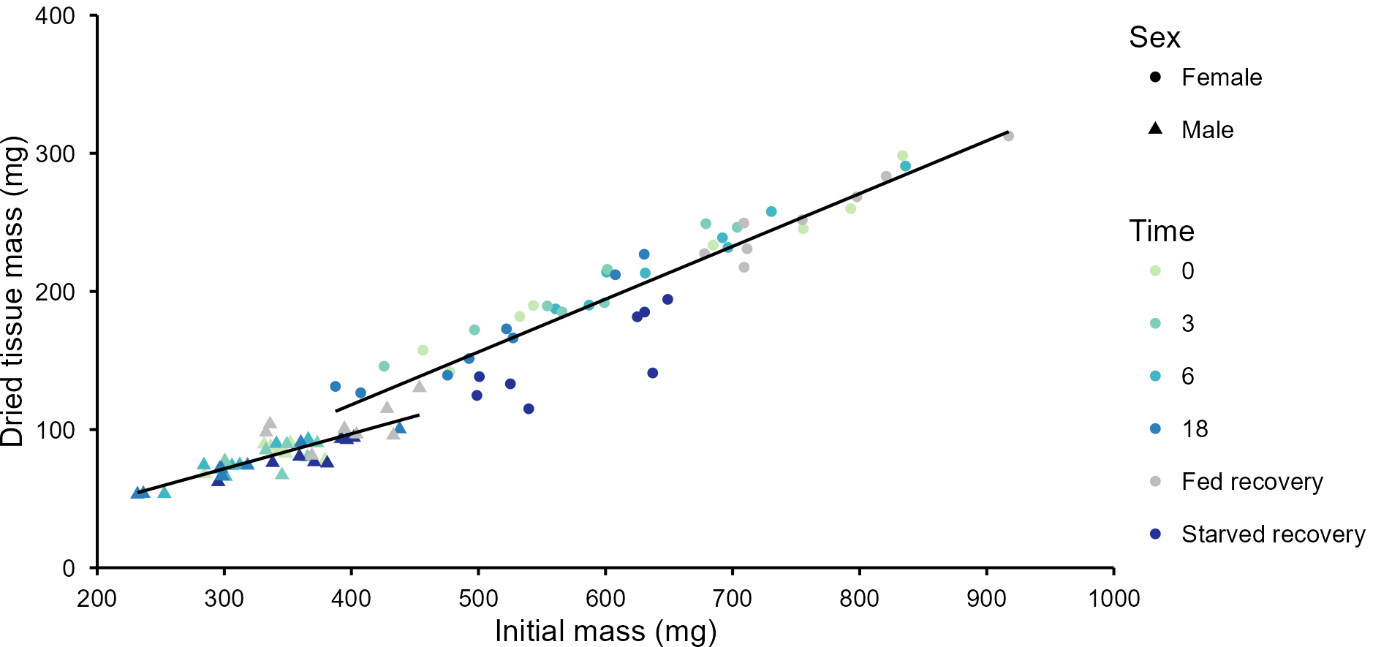

Figure S3. Dried tissue mass (mg) is strongly correlated with initial mass (mg) in Acheta domesticus (n = 48 males and 48 females). R^2^ = 95.4 %.

Table S2. Analysis of Variance table for the reduced model for tissue dry mass. Significant terms are bolded.

|  | df | Sum of Squares | Mean  Sum of Squares | F | P |
| --- | --- | --- | --- | --- | --- |
| Initial mass | 1 | 454322 | 454322 | 4768.096 | **<0.001** |
| Time treatment | 5 | 10428 | 2086 | 21.8878 | **<0.001** |
| Sex | 1 | 2057 | 2057 | 21.5849 | **<0.001** |
| Initial mass:Sex | 1 | 485 | 485 | 5.0907 | **0.02672** |
| Time treatment:Sex | 5 | 4150 | 830 | 8.7114 | **<0.001** |
| Residuals | 82 | 7813 | 95 |  |  |

Table S3. Type III Analysis of Variance Table with Satterthwaite's method for reduced total lipid model. Significant terms are bolded.

|  | Sum of Squares | Mean  Sum of Squares | df | Residual df | F | P |
| --- | --- | --- | --- | --- | --- | --- |
| Initial mass | 229.464 | 229.464 | 1 | 90.73 | 63.4406 | **<0.001** |
| Time treatment | 49.925 | 9.985 | 5 | 92.218 | 2.7606 | **0.02275** |
| Sex | 20.157 | 20.157 | 1 | 92.33 | 5.5728 | **0.02035** |
| Initial mass:Time treatment | 50.316 | 10.063 | 5 | 90.789 | 2.7822 | **0.02199** |
| Initial mass:Sex | 19.188 | 19.188 | 1 | 91.767 | 5.3049 | **0.02352** |
| Time treatment:Sex | 45.67 | 9.134 | 5 | 91.918 | 2.5253 | **0.0345** |

Table S4. Analysis of Variance table for the reduced total protein model. Significant terms are bolded.

|  | df | Sum of Squares | Mean  Sum of Squares | F | P |
| --- | --- | --- | --- | --- | --- |
| Initial mass | 1 | 43410 | 43410 | 643.5666 | **<0.001** |
| Time treatment | 5 | 2356 | 471 | 6.9869 | **<0.001** |
| Sex | 1 | 1964 | 1964 | 29.1209 | **<0.001** |
| Initial mass:Time treatment | 5 | 342 | 68 | 1.0151 | 0.414211 |
| Initial mass:Sex | 1 | 727 | 727 | 10.7759 | **0.001512** |
| Residuals | 82 | 5531 | 67 |  |  |

Table S5. Type III Analysis of Variance Table with Satterthwaite's method for reduced total glycogen model. Significant terms are bolded.

|  | Sum of Squares | Mean Sum of Squares | df | Residual df | F | P |
| --- | --- | --- | --- | --- | --- | --- |
| Initial mass | 6.6012 | 6.6012 | 1 | 90.389 | 19.7891 | **<0.001** |
| Time treatment | 9.2969 | 1.8594 | 5 | 89.825 | 5.574 | **0.000162** |
| Sex | 1.7254 | 1.7254 | 1 | 92.7 | 5.1723 | **0.025257** |
| Initial mass:Sex | 2.384 | 2.384 | 1 | 92.009 | 7.1468 | **0.008885** |
